# Loss of Zbtb20 disrupts cochlear supporting cell differentiation and maturation and extends the postnatal hair cell regenerative window in mice

**DOI:** 10.64898/2026.08.19.745776

**Authors:** Charles Morgan, Zia Ur Rehman, Angelika Doetzlhofer

**Author notes:** Department of Otolaryngology-Head & Neck Surgery, University of Texas Medical Branch, Galveston, TX, 77555, USA.

## Abstract

Cochlear hair cell (HC) loss is a leading cause of hearing loss in humans. HCs can be generated from adjacent supporting cells (SCs); however, this regenerative capacity is lost after the onset of hearing. Using *Emx2*^Cre^ *Zbtb20* knockout mice, we show that ZBTB20 deficiency delays cell-cycle exit, differentiation, and maturation of cochlear SCs. Transcriptomic analysis of postnatal cochlear sensory epithelia indicates that ZBTB20 loss postpones the downregulation of progenitor genes, including *Sox11* and *Hmga2*, and delays activation of a maturation-specific gene program. Additionally, experiments with cochlear organoid and organotypic explant models reveal that prolonged, and to a lesser extent acute, ZBTB20 loss increases the mitotic and HC-regenerative potential of cochlear SCs. Transcriptomic profiling shows that acute ZBTB20 ablation upregulates the midkine receptor *Ptprz1*, and further studies show that exogenous midkine, similar to ZBTB20 loss, promotes cell-cycle reentry and proliferation in cochlear organoid cultures.

## INTRODUCTION

The mammalian auditory sensory epithelium, situated within the cochlea of the inner ear, is essential for auditory perception. This spiral-shaped epithelium is comprised of two types of mechano-sensory hair cells (HCs), termed inner and outer HCs, that are innervated by afferent and efferent neurons and surrounded by distinct supporting cell (SC) subtypes. HCs are responsible for the conversion of sound waves into electrical signals, whereas SCs are required for the development, survival, and function of HCs. Deficiencies in the biophysical properties or function of SCs have been demonstrated to cause auditory dysfunction and hearing loss (Chen et al., 2021; Chrysostomou et al., 2020; Zhu et al., 2013). In non-mammalian vertebrates, such as birds and fish, SCs possess the capacity to regenerate HCs (Brignull et al., 2009). HCs and SCs share a common lineage, both originating from a pool of progenitor cells known as prosensory cells. Following auditory HC loss, SCs in fish and birds acquire progenitor-like characteristics that enable them to form new HCs through mitotic and non-mitotic mechanisms (Jimenez et al., 2022; Takeuchi et al., 2026). In neonatal mice, a small subset of SCs has been observed to re-enter the cell cycle and generate new HCs after HC injury (Bramhall et al., 2014; Cox et al., 2014). However, in mice, HCs and SCs are not functional at birth and require postnatal maturation, which involves changes in cell shape, refinement of intercellular junctions and cytoskeletal organization, and the acquisition of specialized mechanical and homeostatic functions (Ceriani et al., 2025; Walters and Zuo, 2013). The onset of cochlear SC maturation, which in mice occurs at the end of the first week after birth, coincides with their loss of regenerative capacity (Li and Doetzlhofer, 2020; Maass et al., 2015; Tao et al., 2021). However, the extent to which maturation restricts SC competence to respond to regenerative cues, both in vitro and in vivo, remains incompletely understood, and the genes and signaling pathways governing cochlear SC maturation are not well characterized.

A potential candidate for controlling the maturation and regenerative plasticity of cochlear SCs is the transcription factor ZBTB20. ZBTB20 is highly expressed in cochlear SCs during maturation (Xie et al., 2023) but is rapidly downregulated following LIN28B- and TRIM71-mediated reprogramming of cochlear SCs into progenitor-like cells (Li et al., 2022; Li et al., 2023). TRIM71 and LIN28B are evolutionarily conserved RNA-binding proteins that promote stemness and growth (Rehfeld et al., 2015; Worringer et al., 2014). In the developing cochlea, LIN28B and TRIM71 inhibit differentiation by maintaining HC and SC progenitors in a proliferative, undifferentiated state (Golden et al., 2015; Li et al., 2025). At later, postnatal stages, reactivation of these factors has been shown to enhance HC regeneration (Li and Doetzlhofer, 2020; Li et al., 2022; Li et al., 2023). Mutations in *ZBTB20* cause Primrose Syndrome, a rare progressive genetic disorder that can present with sensorineural hearing loss (Cordeddu et al., 2014). A recent study using early otic-stage knockout of *Zbtb20* (Foxg1^Cre^) demonstrated that loss of *Zbtb20* arrests cochlear maturation and causes deafness, which the authors attributed to the absence of root cells adjacent to the lateral edge of the cochlear sensory epithelium (Xie et al., 2023). However, ZBTB20’s potential role in cochlear SC maturation and plasticity is currently unknown.

Using Emx2^Cre^ *Zbtb20* knockout mice, we find that embryonic ablation of ZBTB20 in the developing cochlear duct delays terminal mitosis, differentiation, and maturation of cochlear SCs, causing patterning defects in both the SC and HC layers. In the absence of ZBTB20, postnatal cochlear SCs fail to downregulate progenitor genes or initiate maturation-specific gene expression, with this defect being most pronounced in Hensen cells. Chronic deletion and, to a lesser extent, acute deletion of *Zbtb20* increases the mitotic and HC regenerative potential of postnatal cochlear SCs in vitro. Furthermore, transcriptomic and functional analyses identify Ptprz1-midkine signaling as a pathway through which ZBTB20 restricts the mitotic capacity of cochlear SCs.

## RESULTS

### *Zbtb20* knockout disrupts cell cycle exit, patterning, and differentiation of cochlear SC progenitors

To address the role of ZBTB20 in the embryonic cochlea, we employed a conditional knockout (KO) approach using *Zbtb20* floxed (*Zbtb20^f/f^*) mice, which contain loxP sites flanking a critical exon (de Araújo et al., 2021). These mice were crossed with *Emx2-Cre* (*Emx2^cre/+^*) mice, allowing selective deletion of Zbtb20 in cochlear epithelial cells, including prosensory cells, around E11.5-E12.0 (Kimura et al., 2005). This strategy allowed for the generation of control (*Zbtb20^f/f^; Emx2^+/+^*or *Zbtb20^f/+^; Emx2^+/+^*), heterozygote (*Zbtb20^f/+^; Emx2^cre/+^*), and KO (*Zbtb20^f/f^; Emx2^cre/+^*) conditions. We validated the knockout efficacy at P5 using anti-ZBTB20 immunostaining and RT-qPCR analysis of *Zbtb20* mRNA expression in isolated sensory epithelia (Fig. 1*A, B*). We also confirmed that the knockout mice exhibited severely elevated ABR thresholds in response to broadband and frequency-specific stimuli, as previously reported (Fig. 1*C*) (Xie et al., 2023).

**Fig. 1.**
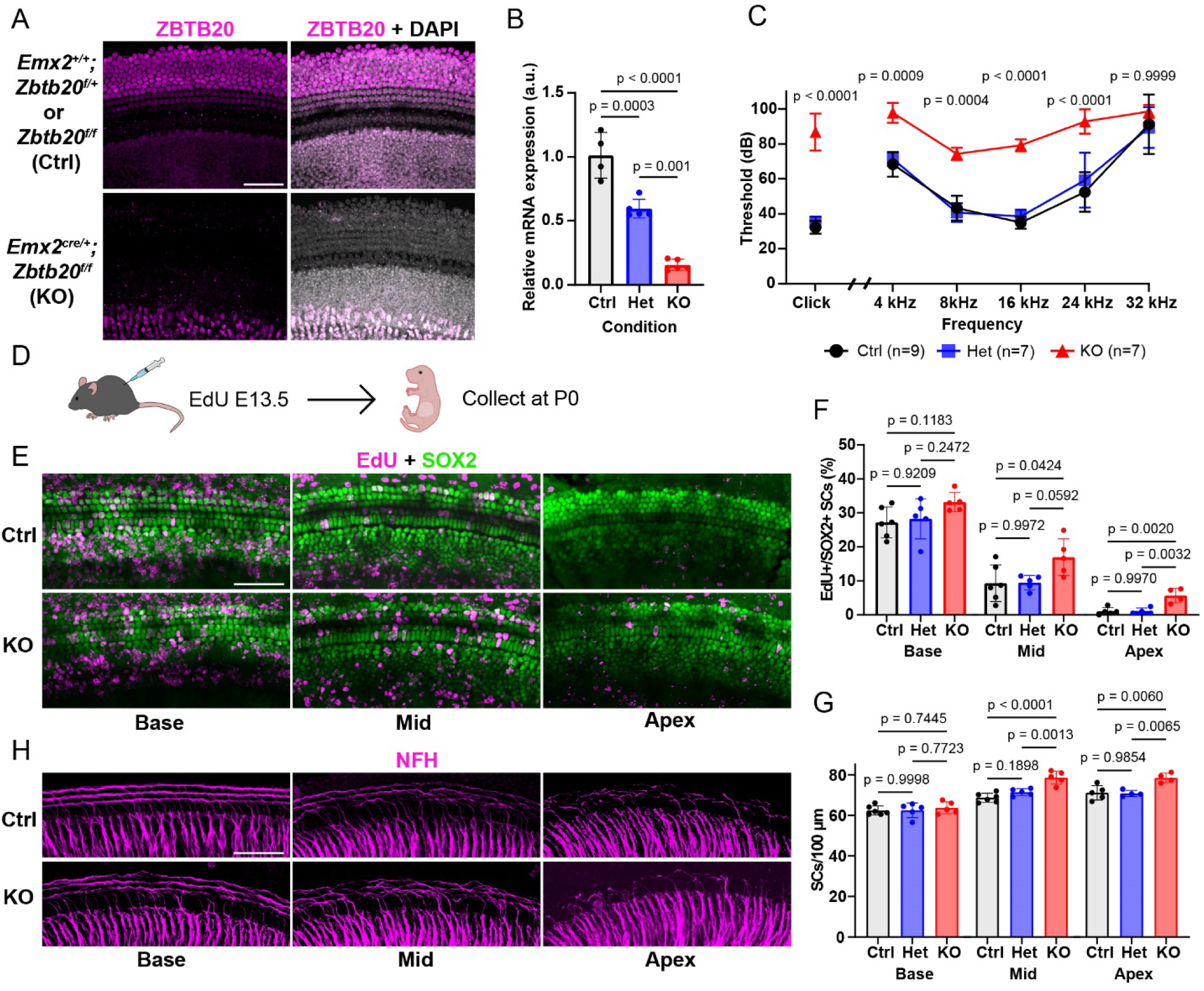
Loss of *Zbtb20* delays cell cycle withdrawal and differentiation of cochlear SC progenitors. (A) Representative confocal images showing immunostaining of ZBTB20 in P5 control and *Emx2^cre/+^;Zbtb20^f/f^*(KO) cochlear sensory epithelia. Scale bar = 50μm. (B) RT-qPCR of *Zbtb20* in isolated P5 cochlear sensory epithelia of control, het, and KO mice. *P* values were determined using a one-way ANOVA and post-hoc Tukey’s HSD test for multiple comparisons. (C) Graph showing ABR thresholds in response to broadband noise (Click) and at multiple frequency levels for control, het, and KO mice. *P* values were determined using a one-way ANOVA and post-hoc Tukey’s HSD test for multiple comparisons. (D) Experimental schematic. Timed-pregnant female mice received a single EdU pulse at E13.5, and cochleae were collected for analysis from control, het, and KO littermates at P0. (E) Representative confocal images of EdU incorporation (magenta) in SCs (SOX2, green) at the base, mid, and apex of P0 control and KO cochlear sensory epithelia. Control, het, and KO mice were collected. Scale bar = 50μm. (F) Quantification of EdU incorporation in SCs as shown in (E). *P* values were determined using a one-way ANOVA and post-hoc Tukey’s HSD test for multiple comparisons. (G) Quantification of SOX2^+^ SCs as shown in (E). *P* values were determined using a one-way ANOVA and post-hoc Tukey’s HSD test for multiple comparisons. (H) Representative confocal images showing immunostaining of neurofilament H (NFH, magenta) at the base, mid, and apex of P0 control and KO cochleae. Control, het, and KO mice were collected. Scale bar = 50μm.

Cochlear HCs and SCs derive from a common pool of SOX2-expressing progenitor cells, termed prosensory cells, that reside at the floor of the developing cochlear epithelial duct. In mice, cochlear prosensory cells exit the cell cycle between embryonic day 12.5 (E12.5) and E14.5 following an apical-to-basal gradient (Chen and Segil, 1999). Re-analysis of previously published single-cell transcriptomic data (Kolla et al., 2020) revealed robust *Zbtb20* mRNA expression in cochlear prosensory cells (Fig. S1, E14). To determine whether *Zbtb20* KO alters terminal mitosis of prosensory cells, we administered a pulse of EdU at E13.5 and analyzed EdU incorporation in cochlear HCs and SCs (Deiter’s cells and pillar cells) in newborn *Zbtb20* KO, heterozygote (Het), and control mice [postnatal day 0 (P0)](Fig. 1D). Anti-MYO7A and SOX2 immunostaining was used to identify HCs and SCs, respectively. We found that EdU labeled a significantly higher percentage of SCs in the mid and apical turns of KO mice compared to control mice (Fig. 1*E*, *F*). Additionally, SOX2 immunostaining showed an increase in the total number of SCs in the mid and apical turns of KO mice compared to control mice, as well as a breakdown in the uniform three rows of Deiter’s cells observed in control tissue (Fig. 1*G*). Interestingly, the percentage of MYO7A^+^ EdU^+^ HCs in *Zbtb20* KO mice was similar to that of control mice, and no changes in the total number of IHC or OHCs were observed (Fig. S2*A-G*). At around E14.5, prosensory cells in the mid-base begin to differentiate following a steep base-to-apex gradient that reaches the apex at around E17/E18 (Chen et al., 2002). Potential defects in HC and SC differentiation can be assessed by examining the HC bundle (stereocilia) morphology and by examining the innervation pattern of type II spiral ganglion neurons (SGNs) in the lateral compartment of the sensory epithelium (Coate and Kelley, 2013; Ghimire and Deans, 2019). At P0, HCs in the apex don’t have well-formed stereocilia. While OHCs in the basal turn are fully innervated by type II SGNs, forming three rows of fibers, OHCs in the apical turn are not innervated, with the axons having just initiated turning. In P0 *Zbtb20* KO mice, we observed that this innervation pattern was delayed relative to control littermates (Fig. 1*H*). This delay, though slight, was observed at all three turns of KO cochleae. By contrast, knockout of *Zbtb20* did not delay cochlear HC differentiation, as judged by the similar morphology of HC stereocilia in P0 *Zbtb20* KO mice and control littermates (Fig. S2*H*). This decoupling of HC and SC differentiation is highly unusual but has recently been reported in mice, in which Deiters cell/pillar cell transcription factor PROX1 was knocked out (Kale et al., 2025). Together, these data indicate that ZBTB20 plays an essential role in the terminal differentiation of cochlear SCs.

### *Zbtb20* knockout results in ectopic postnatal HCs and SCs

To further evaluate the role of ZBTB20 in cochlear sensory patterning, we examined the cochlear sensory epithelia of control and KO mice 5 days later (P5). Immunostaining for MYO7A and PROX1, which selectively mark Deiter’s cells and pillar cells, revealed extra OHCs and Deiter’s cells (Fig. 2*A-C*). Cells were considered to be ectopic if they were supernumerary to the three rows of OHCs and Deiter’s cells found in control tissue. In contrast to P0, significantly more ectopic OHCs were observed in all regions of KO cochleae at P5 compared to control samples (Fig. 2*B*). This result was confirmed by counting the total number of OHCs. In contrast, the total number of IHCs was only significantly increased in the mid-turn of KO cochleae (Fig. S3*A, B*). As with OHCs, significantly more ectopic PROX1^+^ SCs (specifically Deiter’s cells) were identified in all regions of KO cochleae compared to control samples (Fig. 2*C*).

**Fig. 2.**
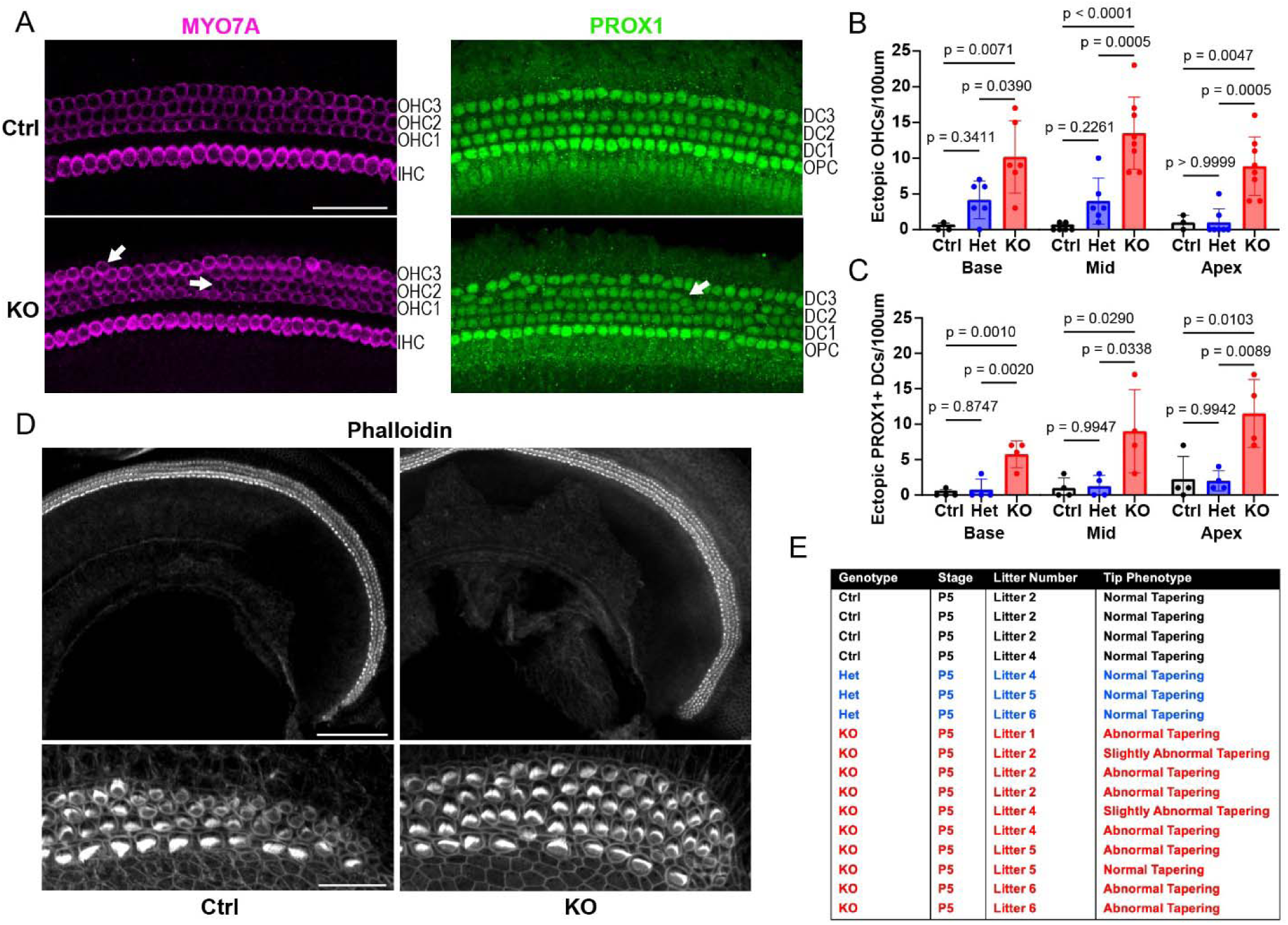
Loss of *Zbtb20* leads to ectopic cell production and patterning defects in the lateral cochlear sensory epithelium, especially at the apical tip. (A) Representative confocal images showing immunostaining of HCs (MYO7A, magenta) and SCs (SOX2, green) at the mid-turn of P5 control and KO cochlear sensory epithelia. Control, het, and KO mice were collected. Scale bar = 50μm. (B) Quantification of ectopic OHCs as shown in (A). Ectopic cells were defined as those that are located outside of the conventional arrangement of three rows. *P* values were determined using a one-way ANOVA and post-hoc Tukey’s HSD test for multiple comparisons. (C) Quantification of ectopic Deiter’s cells as shown in (A). Ectopic cells were defined as those that are located outside of the conventional arrangement of three rows. *P* values were determined using a one-way ANOVA and post-hoc Tukey’s HSD test for multiple comparisons. (D) Representative confocal images showing phalloidin staining of F-actin in P5 control and KO cochlear sensory epithelia. Control, het, and KO mice were collected. Upper panels show the apical turn at 10X magnification, whereas lower panels show the apical tip at 63X magnification. Scale bars = 100μm (upper panel) and 20μm (lower panel). (E) Chart showing the penetrance of the observed apical tip phenotype in (D).

In addition to instances of ectopic HCs and SCs across the length of the cochlear sensory epithelium of KO mice, we identified a striking phenotype at the apical tip (Fig. 2*D*). In control tissue, tapering is observed at the apical tip, mediated by a decrease in OHC number and reduction of rows from three down to one. In the corresponding *Zbtb20* KO tissue, we observed no tapering, with an additional row of OHCs compared to the expected reduction. This phenotype was highly penetrant, as it was observed in nearly all KO samples (Fig. 2*E*). SOX2 is transiently expressed in newly formed HCs but absent from cochlear HCs at later stages such as P5 (Atkinson et al., 2018). Consistent with previous findings, at P5, IHCs and OHCs in control mice lacked SOX2 expression. In corresponding *Zbtb20* KO mice, we detected a few SOX2-expressing IHCs (Fig. S3*C*). EdU pulses administered from P0 to P5 revealed essentially no labeling of HCs in either control or KO tissue, with only a few apical IHCs found labeled with EdU (Fig. S3*D, E*). These data indicate that postnatal HCs are added via a non-mitotic process.

### Induction of maturation-specific gene programs is delayed in *Zbtb20-*deficient cochlear sensory epithelia

Following differentiation, *Zbtb20* expression is downregulated in IHCs and OHCs but remains highly expressed in SCs as they begin to mature (Fig. S1, P0 and P7). To gain insight into the molecular and cellular mechanisms through which ZBTB20 controls cochlear SC maturation, we analyzed gene expression in cochlear sensory epithelia isolated from P5 control, heterozygous, and *Zbtb20* KO mice using RNA sequencing (RNA-seq). Analysis of the obtained RNA-seq data revealed 676 significantly differentially expressed genes (DEGs) using an adjusted p-value (q-value) threshold of <0.05 (Fig. 3*A* and S4)(Table S1). 489 of these DEGs were upregulated, whereas only 187 were downregulated, supporting the role of ZBTB20 as a transcriptional repressor (Xie et al., 2008; Zhang et al., 2015). Among the downregulated genes were SC genes critical for auditory sensory development (e.g., *Gdf6*)(Zafeer et al., 2026) and hearing (e.g., *Aqp4, Gjb6*)(Mhatre et al., 2002; Teubner et al., 2003). Among the upregulated genes were the cochlear progenitor genes *Hmga2* and *Sox11*, and increased expression of these genes in cochlear SCs is linked to increased regenerative plasticity (Gnedeva and Hudspeth, 2015; Li et al., 2023). Gene Ontology (GO) analysis of DEGs indicates that loss of ZBTB20 impairs pathways associated with sensory organ development, ERK1/2 signaling, ion homeostasis and lipid metabolism, while activating pro-growth pathways such as PI3K-Akt-mTOR signaling and wound healing (immune response/inflammation) (Fig. 3*B*). To confirm the findings of our RNA-seq analysis, we analyzed the expression of FABP7 and HMGA2 in P5 control and *Zbtb20* KO cochlear sensory epithelia using immunostaining. Our analysis revealed that ZBTB20 loss differentially affected FABP7 expression in Hensen cells and inner phalangeal cells, with FABP7 expression in Hensen cells nearly absent in the absence of ZBTB20, whereas its expression in inner phalangeal cells remained unchanged compared to control tissue (Fig. 3*C, D*). Immunostaining against HMGA2 protein showed increased expression of HMGA2 in cochlear SCs and lateral non-sensory epithelial cells at the mid and apical turns, in *Zbtb20* KO tissue compared to control tissue (Fig. 3*E*). We next compared our dataset to a publicly available dataset of *Zbtb20* KO whole cochlea at P10 to identify genes that were persistently downregulated or upregulated relative to their respective controls, which would help evaluate the severity and persistence of maturation delay (Xie et al., 2023). There were 126 DEGs that were common to both datasets, including *Fabp7* and *Aqp4,* and hierarchical clustering of these DEGs revealed patterns of co-regulation (Fig. 3*F*).

**Fig. 3.**
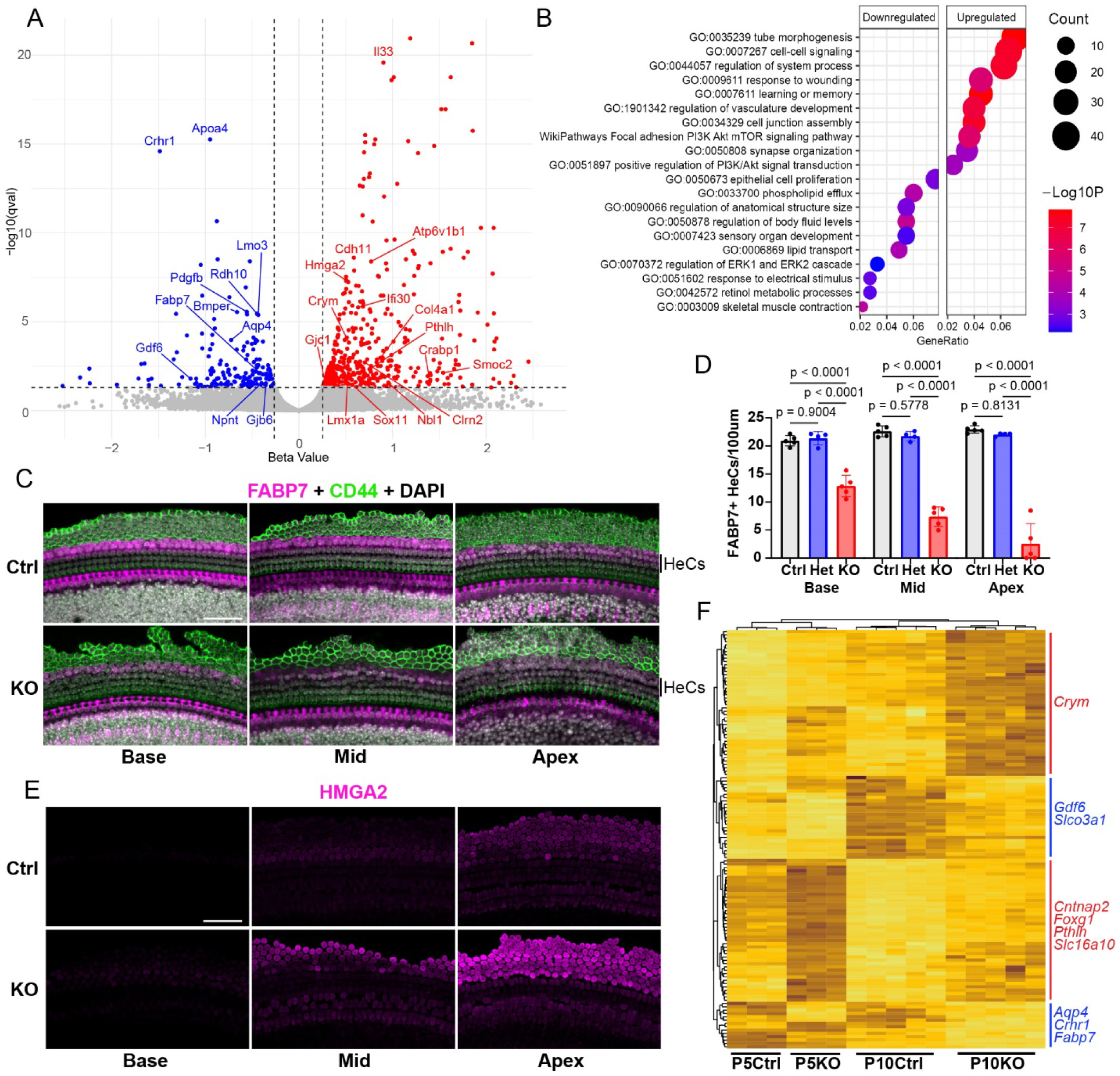
*Zbtb20* KO cochlear sensory epithelia exhibit a delay in cell maturation. (A) Volcano plot of RNA-seq data collected from P5 control and KO cochlear sensory epithelia. Control, het, and KO mice were collected. Individual genes are plotted according to beta-value (*x*-axis) and −log_10_(q-value) (*y*-axis). Blue = gene was downregulated in KO tissue, red = gene was upregulated in KO tissue. Significance of differential expression was determined using the Wald test with Benjamini-Hochberg p-value correction. (B) Gene ontology analysis. Analysis was segregated by direction of regulation. Categories are ranked by GeneRatio, the number of differentially expressed genes in a category divided by the total number of differentially expressed genes. (C) Representative confocal images showing immunostaining of Hensen cells and inner phalangeal cells (FABP7, magenta) as well as outer pillar cells and Claudius cells (CD44, green) at the base, mid, and apex of P5 control and KO cochleae. Control, het, and KO mice were collected. Scale bar = 50μm. (D) Quantification of FABP7 immunostaining in Hensen cells shown in (C). *P* values were determined using a one-way ANOVA and post-hoc Tukey’s HSD test for multiple comparisons. (E) Representative confocal images showing immunostaining of HMGA2 (magenta) at the base, mid, and apex of P5 control and KO cochleae. Control, het, and KO mice were collected, and all images were collected using identical digital gain and laser power settings to highlight the observed basal-apical gradient of protein expression. Scale bar = 50μm. (F) Heatmap of differentially expressed genes in the dataset generated from this study as shown in (A) and of differentially expressed genes in a dataset generated from P10 whole cochleae of control and *Foxg1^cre/+^;Zbtb20^f/f^* (KO) mice. Genes which were found to be differentially expressed in both datasets are shown in middle overlapping field.

### Chronic deletion of *Zbtb20* promotes organoid formation and the HC production capacity of cochlear SCs

Early postnatal cochlear SCs and adjacent Kölliker’s organ cells (KCs) retain the capacity to reenter the cell cycle and form HCs, but their capacity to do so rapidly declines at the end of the first postnatal week when the cochlear sensory epithelium starts to undergo maturation (Maass et al., 2015; Tao et al., 2021). Considering the delayed differentiation and maturation observed in *Zbtb20*-deficient cochlear sensory epithelia in vivo, we assessed whether *Zbtb20*-deficient cochlear SCs (and KCs) exhibit greater proliferative and/or HC regenerative capacity compared to wild-type cochlear SCs (and KCs) using a previously described organoid culture system (McLean et al., 2017). Cochlear sensory epithelia from P5 control, heterozygous and *Zbtb20* KO were isolated and the dissociated cells containing SCs were plated in Matrigel matrix droplets and cultured with epidermal growth factor (EGF), fibroblast growth factor 2 (FGF2), the GSK3B inhibitor CHIR99021, the histone deacetylase inhibitor valproic acid (VPA), and the TGFBR1 inhibitor 616452, which together form a pro-growth expansion media (Fig. 4*A*). We found that cochlear epithelial cells from *Zbtb20* KO mice formed organoids which were on average significantly larger than that from heterozygote or control mice (Fig. 4*B, D*). Additionally, we found that *Zbtb20* KO cells formed organoids at a significantly higher rate (the number of organoids formed as a percentage of the number of cells plated) than heterozygote or control cells (Fig. 4*E*). A two-hour EdU pulse, which labels cells in S phase, revealed that *Zbtb20* KO organoids contained a significantly higher number of actively proliferating cells, which based on their high expression of SC-specific Notch ligand JAG1 constituted SC-like cells (Fig. 4*C, F*).

**Fig. 4.**
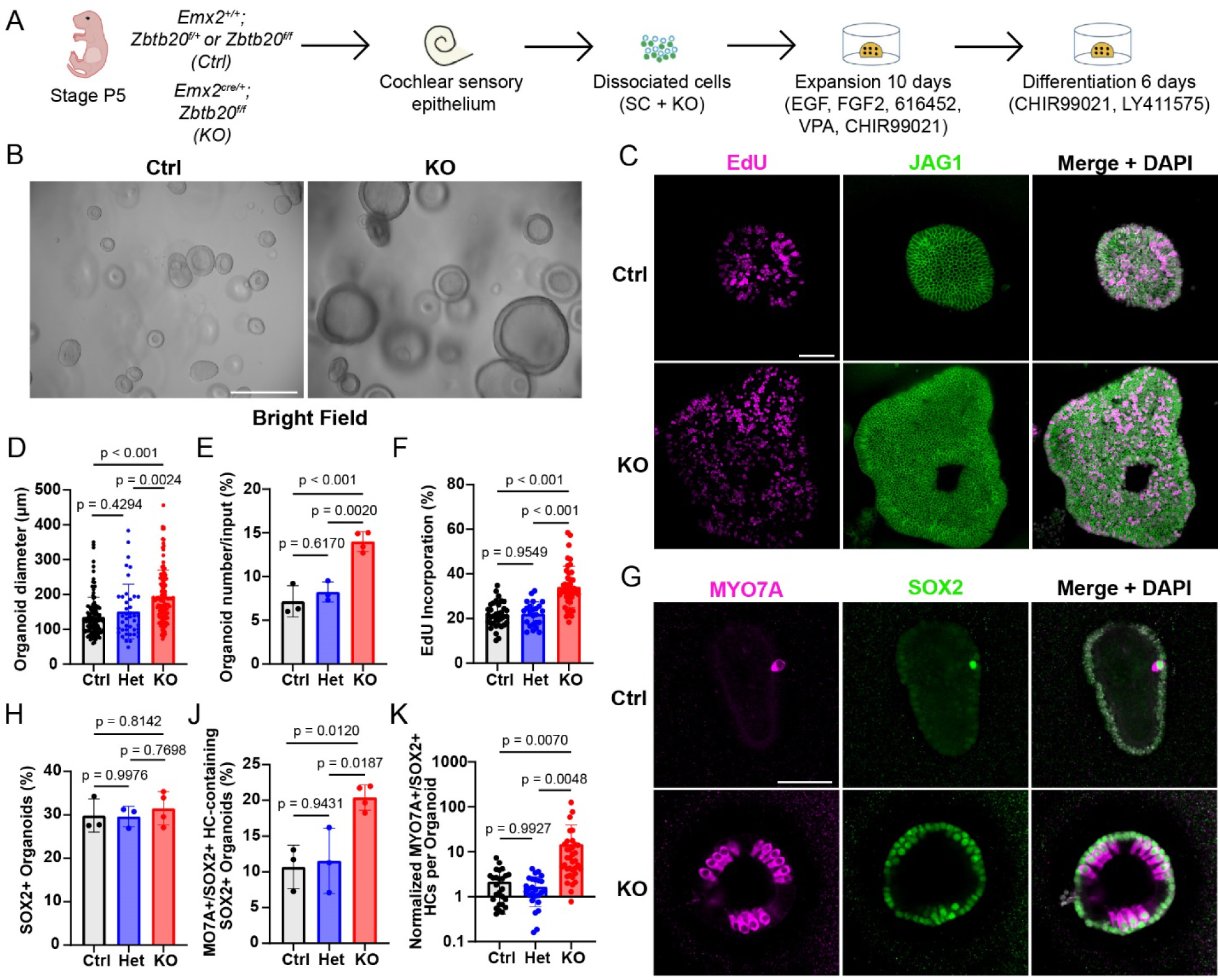
Chronic deletion of *Zbtb20* enhances the proliferative and HC-forming capacity of P5 cochlear SCs. (A) Experimental schematic of organoid generation and HC differentiation. Made with graphics from scidraw.io. (B) Representative bright-field images of control and KO organoid cultures after 10DIV. Scale bar = 200μm. (C) Representative confocal images showing EdU incorporation (magenta) and immunostaining of JAG1 (green) in control and KO organoids after 10DIV. Control, het, and KO organoids were collected. Scale bar = 50μm. (D) Quantification of organoid diameter as shown in (B). *P* values were determined using a one-way ANOVA and post-hoc Tukey’s HSD test for multiple comparisons. (E) Quantification of organoid formation efficiency as shown in (B). Efficiency was defined as the percentage of organoids observed in a culture relative to the number of cells plated. *P* values were determined using a one-way ANOVA and post-hoc Tukey’s HSD test for multiple comparisons. (F) Quantification of EdU incorporation within JAG1^+^ organoids as shown in (C). *P* values were determined using a one-way ANOVA and post-hoc Tukey’s HSD test for multiple comparisons. (G) Representative confocal images showing immunostaining of HCs (MYO7A, magenta) and SOX2 (green) in control and KO organoids after 16DIV. Control, het, and KO organoids were collected. Scale bar = 50μm. (H) Quantification of the percentage of SOX2^+^ organoids. *P* values were determined using a one-way ANOVA and post-hoc Tukey’s HSD test for multiple comparisons. (J) Quantification of the percentage of SOX2^+^ organoids containing at least one MYO7A^+^SOX2^+^ nascent HC as shown in (G). *P* values were determined using a one-way ANOVA and post-hoc Tukey’s HSD test for multiple comparisons. (K) Quantification of MYO7A^+^ SOX2^+^ nascent HCs as shown in (G). Counts were normalized based on organoid area to account for differences in organoid size across conditions. Data are displayed on a Log_10_ scale. *P* values were determined using a one-way ANOVA and post-hoc Tukey’s HSD test for multiple comparisons.

Next, we tested the organoid-forming and proliferative capacities of SCs purified using fluorescence-activated cell sorting (FACS) with the *Lfng*–green fluorescent protein (GFP) transgene (Fig. S5*A, B*). In this model, all conditions express GFP in the majority of cochlear SC subtypes, including Deiters’ cells, outer pillar cells, inner phalangeal cells, and border cells, but not in KCs, thereby allowing evaluation exclusively of cochlear SCs (Korrapati et al., 2013). After 18 days in expansion, we found that *Zbtb20* KO *Lfng*-GFP^+^ cells formed significantly larger organoids than the *Lfng*-GFP^+^ cells derived from control or heterozygote conditions, and that the organoid formation rate was about threefold greater in *Zbtb20* KO cultures than control cultures (Fig. S5*C-E*).

We then evaluated whether ZBTB20 loss enhances HC formation. We switched unsorted, P5 sensory epithelium-derived cultures from the expansion medium to a differentiation medium containing CHIR99021 and Notch inhibitor LY411575 for six days to promote HC formation (Fig. 4*A*). We then immuno-stained organoids for MYO7A and SOX2 expression. Co-expression of HC marker MYO7A and SOX2 marks nascent HCs, while SOX2 expression by itself marks non-sensory cells that are competent to form HCs (SCs and KCs) (Atkinson et al., 2018). Our analysis revealed that ZBTB20 loss did not alter the percentage of organoids that contained SOX2^+^ cells (Fig. 4H), but *Zbtb20* KO organoids produced significantly more HCs than control or heterozygote organoids (Fig. 4*G*). To quantitatively describe the observed result, we first confirmed that cultures from each condition produced a similar percentage of SOX2+ organoids. (Fig. 4*H*). Then, we calculated the percentage of SOX2^+^ organoids that produced at least one MYO7A^+^SOX2^+^ HC and found it to be higher in the *Zbtb20* KO cultures (Fig. 4*J*). Finally, we determined the number of MYO7A^+^SOX2^+^ HCs per SOX2^+^ organoid. Given the significant difference in organoid size between the *Zbtb20* KO and control/heterozygote conditions, we normalized the number of MYO7A^+^SOX2^+^ HCs in an organoid by the size of that organoid. The resulting data show that *Zbtb20* KO organoids produce significantly more HCs per organoid than control or heterozygote organoids (Fig. 4*K*).

### ZBTB20 limits cochlear SC proliferation through targeting Ptprz1-midkine signaling

To determine whether ZBTB20 remains required to restrict SC plasticity, we acutely knocked out *Zbtb20* in cochlear organoids using a doxycycline (dox)-inducible Cre strategy. To this end, we generated stage P5 *Zbtb20^f/f^*mice that also carried the *R26rtTA*M2* and *TetO-Cre* transgenes (KO-ready), and control littermates that lacked TetO-Cre transgene, and cultured their cochlear epithelial cells as organoids in the presence of doxycycline (dox) (Fig. 5*A*). We first confirmed that dox-mediated *Zbtb20* KO was successful by RT-qPCR analysis of *Zbtb20* expression levels (Fig. 5*B*). We found that *Zbtb20* KO cells formed organoids which were on average significantly larger than heterozygote or control cells (Fig. 5*C, D*). We also found that *Zbtb20* KO cells formed organoids at a significantly higher rate than control cells (Fig. 5*E*). These results mirror the phenotype observed in the chronic deletion (*Zbtb20^f/f^; Emx2^cre/+^*) model, but with a smaller difference between the respective KO and control conditions.

**Fig. 5.**
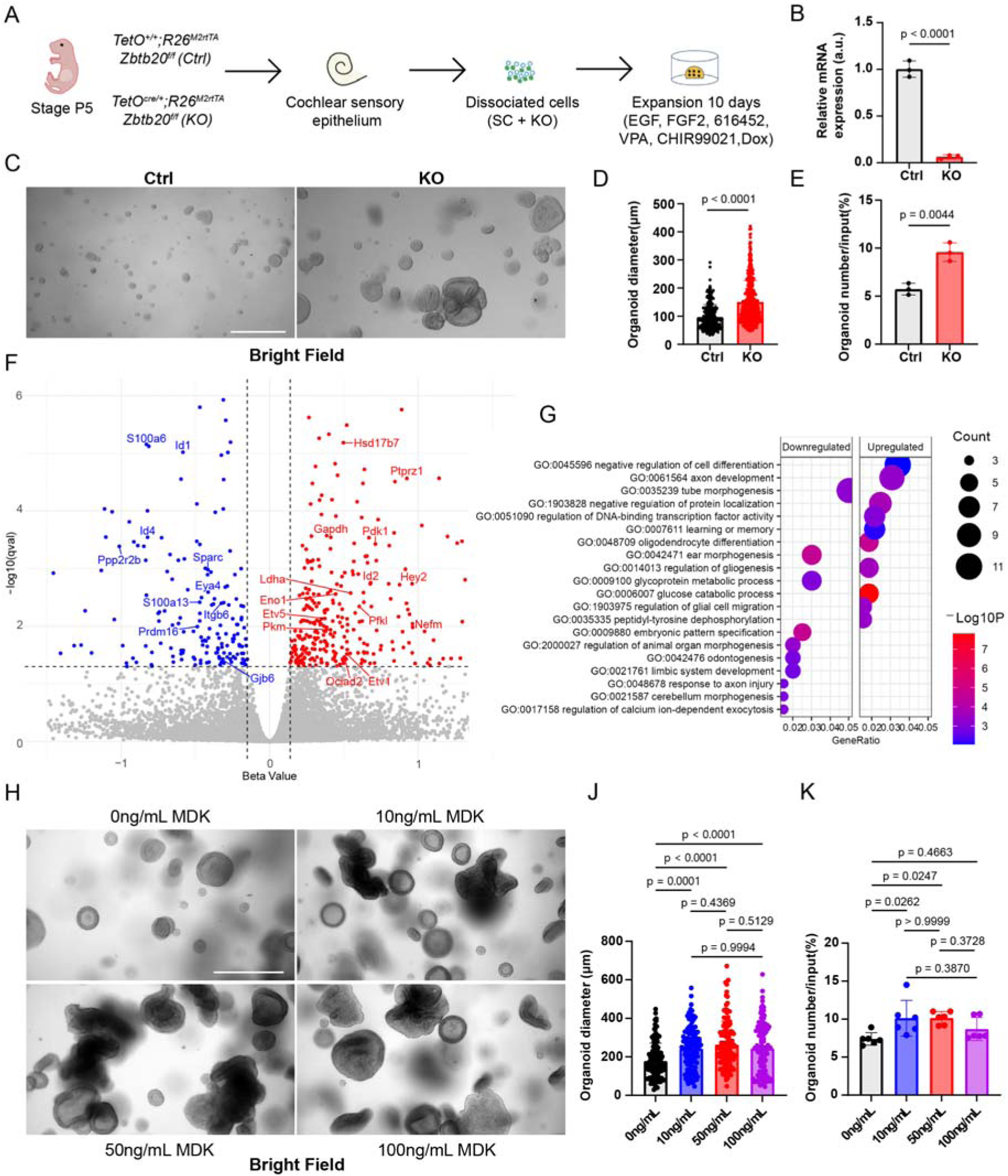
Acute deletion of Zbtb20 enhances cochlear SC proliferation by targeting Ptprz1-midkine signaling. (A) Experimental schematic of organoid generation. Dox was added at plating to induce *Cre* recombination. Made with graphics from scidraw.io. (B) RT-qPCR of *Zbtb20* in control and KO organoid cultures after 10DIV. *P* values were determined using a two-tailed, unpaired t-test. (C) Representative bright-field images of control and KO organoid cultures after 10DIV. Scale bar = 200μm. (D) Quantification of organoid diameter as shown in (B). *P* values were determined using a two-tailed, unpaired t-test. (E) Quantification of organoid formation efficiency as shown in (B). Efficiency was defined as the percentage of organoids observed in a culture relative to the number of cells plated. *P* values were determined using a two-tailed, unpaired t-test. (F) Volcano plot of RNA-seq data collected from control and KO organoids after 10 DIV. Individual genes are plotted according to beta-value (*x*-axis) and −log_10_(q-value) (*y*-axis). Blue = gene was downregulated in KO tissue, red = gene was upregulated in KO tissue. Significance of differential expression was determined using the Wald test with Benjamini-Hochberg p-value correction. (G) Gene ontology analysis. Analysis was segregated by direction of regulation. Categories are ranked by GeneRatio, the number of differentially expressed genes in a category divided by the total number of differentially expressed genes. (H) Representative bright-field images of control organoid cultures after 10DIV. Cultures were supplemented with 0, 10, 50, or 100 ng/mL of midkine (MDK) for the duration of the experiment. Scale bar = 200μm. (J) Quantification of organoid diameter as shown in (H). *P* values were determined using a one-way ANOVA and post-hoc Tukey’s HSD test for multiple comparisons. (K) Quantification of organoid formation efficiency as shown in (J). Efficiency was defined as the percentage of organoids observed in a culture relative to the number of cells plated. *P* values were determined using a one-way ANOVA and post-hoc Tukey’s HSD test for multiple comparisons.

To gain insights into how ZBTB20 may repress cochlear SC re-entry and proliferation, we profiled gene expression in control and *Zbtb20* KO cultures during the expansion phase. Analysis of the RNA-seq data revealed 524 DEGs using a q-value threshold of <0.05 (Fig. 5*F*)(Table S2). As observed in the RNA-seq dataset collected from P5 epithelia of control and chronic *Zbtb20* KO conditions, more DEGs were upregulated (325) than downregulated (199). Many of the downregulated genes were highly expressed in SCs and/or KCs and had essential functions in cell-cell communication (e.g., *Gjb6*)(Jagger and Forge, 2015), extracellular matrix remodeling (e.g., *Sparc*)(Rotllant et al., 2008), and SC/KC differentiation/maturation (*e.g., Prdm16*)(Ebeid et al., 2022). Among the upregulated genes were those involved in glycolysis (*Gapdh, Ldha*, *Pkm*, *Eno1*)(Dienel, 2019). Gene ontology analysis of upregulated and downregulated DEGs identified categories such as negative regulation of cell differentiation, regulation of DNA-binding transcription factor activity, and ear morphogenesis (Fig. 5*G*).

One gene of interest, which was differentially upregulated, *Ptprz1*, encodes a receptor-type protein tyrosine phosphatase. Together with the ligand Midkine (MDK), it comprises a signaling pathway that regulates tyrosine phosphorylation and has been shown to influence cell survival and proliferation (Le et al., 2025; Xia et al., 2019). We tested the hypothesis that the MDK-PTPRZ1 signaling axis regulates SC organoid formation and growth by adding exogenous MDK (0, 10, 50, or 100 ng/mL) to the expansion media (Fig. 5*H*). Profiling of organoid diameter revealed a significant increase in organoid size between all concentrations of MDK and control organoids (Fig. 5*J*). Interestingly, we found that MDK-treated SCs formed organoids at a significantly higher rate than untreated cells at two concentrations, 10 and 50 ng/mL, but not at 100ng/mL, implying a dose-dependent role of MDK-PTPRZ1 signaling in SC survival and organoid formation (Fig. 5*K*).

### Chronic and acute Zbtb20 deletion differentially affect HC regeneration in damaged and undamaged cochlear explants

Although the organoid culture system is advantageous for its high sensitivity in detecting small changes in SC plasticity, it lacks the tissue architecture and microenvironment of the cochlear sensory epithelium. Thus, we decided to test the HC-forming capacity of *Zbtb20* KO SCs in cochlear tissue explants (Fig. 6*A*). Additionally, we employed a genetic strategy to selectively ablate HCs, in which mice express the human diphtheria toxin receptor (DTR) under the control of HC-specific gene *Pou4f3* (*Pou4f3^DTR/+^*) (Golub et al., 2012). This system promotes the death of HCs when diphtheria toxin (DT) is added to culture media, allowing us to study the HC-forming capacity of *Zbtb20* KO SC’s in both damaged (DTR+) and undamaged (DTR-) paradigms. Explants were cultured in DT-containing medium for 1 day, followed by a washout to remove DT, and then cultured in CHIR99021-containing medium (Fig. 6*B*). After 3 days *in vitro*, LY411575 was added to further promote nascent HC differentiation. We then evaluated nascent HC production by immunostaining for MYO7A and SOX2. We found that in undamaged (DTR-) explants, significantly more HCs were produced in the basal and mid turns of the cochlear explants in *Zbtb20* KO compared to control and heterozygote conditions (Fig. 6*C, D*). Similarly, we found that in damaged (DTR+) explants, significantly more HCs were produced in the basal and mid turns of the cochlear explants in *Zbtb20* KO compared to control and heterozygote conditions (Fig. 6*E, F*).

**Fig. 6.**
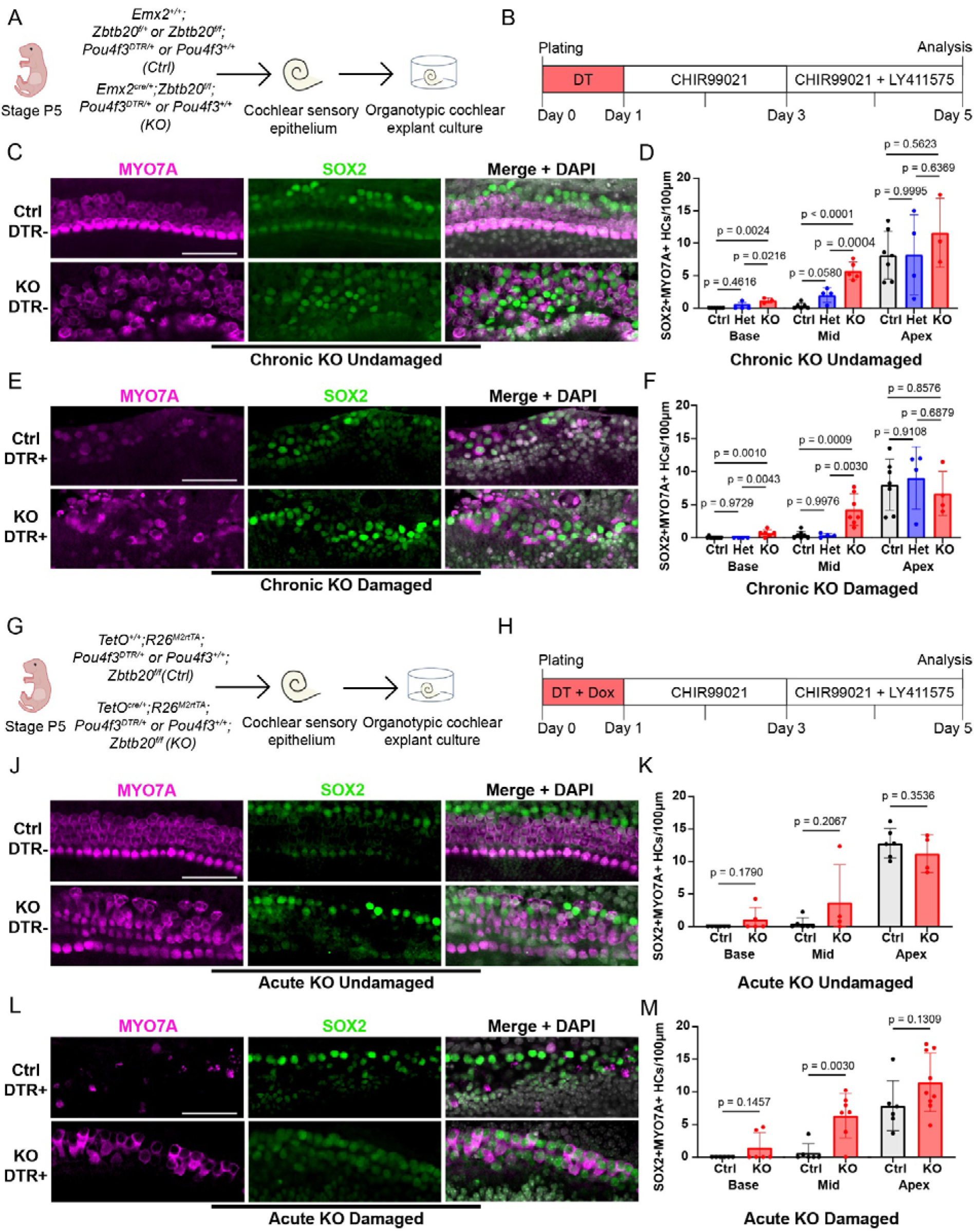
Chronic and acute deletion of *Zbtb20* variably influence the capacity of SCs to transdifferentiate into HCs in damaged and undamaged P5 cochlear sensory epithelia. (A) Experimental schematic of cochlear explant generation using *Emx2^cre/+^*(chronic deletion) mice. Made with graphics from scidraw.io. (B) Time course of DT-mediated HC ablation and HC formation. (C) Representative confocal images showing immunostaining of HCs (MYO7A, magenta) and SOX2 (green) in control and chronic KO undamaged (*Pou4f3^+/+^*) explant cultures after 5DIV. Control, het, and KO explant cultures were collected. Scale bar = 50μm. (D) Quantification of MYO7A^+^SOX2^+^ nascent HCs as shown in (C). *P* values were determined using a one-way ANOVA and post-hoc Tukey’s HSD test for multiple comparisons. (E) Representative confocal images showing immunostaining of HCs (MYO7A, magenta) and SOX2 (green) in control and chronic KO-damaged (*Pou4f3^DTR/+^*) explant cultures after 5DIV. Control, het, and KO explant cultures were collected. Scale bar = 50μm. (F) Quantification of MYO7A^+^ SOX2^+^ nascent HCs as shown in (E). *P* values were determined using a one-way ANOVA and post-hoc Tukey’s HSD test for multiple comparisons. (G) Experimental schematic of cochlear explant generation using *TetO^cre/+^*(acute deletion) mice. Made with graphics from scidraw.io. (H) Time course of DT-mediated HC ablation and HC formation. Dox was added at plating to induce *Cre* recombination. (J) Representative confocal images showing immunostaining of HCs (MYO7A, magenta) and SOX2 (green) in control and acute KO undamaged (*Pou4f3^+/+^*) explant cultures after 5DIV. Control, het, and KO explant cultures were collected. Scale bar = 50μm. (K) Quantification of MYO7A^+^ SOX2^+^ nascent HCs as shown in (C). *P* values were determined using a two-tailed, unpaired t-test. (L) Representative confocal images showing immunostaining of HCs (MYO7A, magenta) and SOX2 (green) in control and acute KO damaged (*Pou4f3^DTR/+^*) explant cultures after 5DIV. Control, het, and KO explant cultures were collected. Scale bar = 50μm. (M) Quantification of MYO7A^+^ SOX2^+^ nascent HCs as shown in (E). *P* values were determined using a two-tailed, unpaired t-test.

We next investigated whether acute deletion of *Zbtb20* at the time of culture establishment would be sufficient to drive increased HC production. Control (*TetO^+/+^; R26^M2rtTA^;Zbtb20^f/f^*) and KO (*TetO^Cre/+^;R26^M2rtTA^;Zbtb20^f/f^*) SCs were profiled using the same explant system, culture parameters (with the addition of dox at plating), and damaged paradigm as described above (Fig. 6*G, H*). Immunostaining for MYO7A and SOX2 revealed that in undamaged (DTR-) explants, acute deletion of *Zbtb20* was not sufficient to drive HC production in any region of the cochlear explants (Fig. 6*J, K*). Interestingly, we found that in damaged (DTR) explants, acute deletion of *Zbtb20* did drive HC production in the mid-turn of the cochlear explants (Fig. 6*L, M*). These data provide evidence of a continued requirement for ZBTB20-mediated repression for enforcing SC identity and limiting HC regenerative plasticity of postnatal SCs.

## DISCUSSION

Cochlear HCs are essential for auditory function, and millions of individuals worldwide experience age-related or noise-induced hearing loss due to HC loss. Recent investigations of the murine cochlear sensory epithelium have demonstrated that SCs possess an age-dependent regenerative capacity to form HCs. This capacity is evident in response to injury, modulation of signaling pathways, or genetic perturbations during early postnatal development, but is lost in the fully mature cochlea. Increasing evidence implicates cell-autonomous maturation processes, including alterations in DNA methylation, chromatin accessibility, and gene regulatory networks, in the decline of SC regenerative plasticity. This study identifies a role for ZBTB20 in regulating cell-autonomous mechanisms that contribute to the loss of SC regenerative capacity during maturation. Analysis of *Emx2^Cre^ Zbtb20* knockout mice reveals that ZBTB20 is necessary for terminal mitosis, differentiation, and maturation of cochlear SCs. Loss of ZBTB20 activates genes and signaling pathways associated with stemness and proliferative potential, thereby prolonging the period during which SC-to-HC regeneration can occur.

The role of ZBTB20 in cell proliferation is multifaceted (Stoyanov et al., 2023). Depending on the specific cell type or tissue, ZBTB20 has been reported to either promote (Zhang et al., 2018) or inhibit cell proliferation (Cao et al., 2016). Here, we uncover that ZBTB20 selectively represses cell division within the SC lineage. We show that deletion of *Zbtb20* selectively delays the cell cycle exit of prosensory cells that give rise to SCs. Such SC lineage-specific delay in terminal mitosis is atypical. In the mammalian cochlea, cell-cycle withdrawal of prosensory cells (HC and SC progenitors) is controlled by the cell-cycle inhibitor p27/Kip1(Chen and Segil, 1999). However, once withdrawn from the cell cycle, SCs, but not HCs, can reenter the cell cycle, requiring continued reinforcement of their post-mitotic state (Jahanshir et al., 2025). These findings suggest that the increased percentage of proliferating SC progenitor cells observed in E13.5 *Zbtb20* knockout mice is due to cell-cycle reentry rather than a delay in the initial cell-cycle exit. This interpretation would imply an early role for ZBTB20 in promoting SC identity through SC-specific gene regulatory programs. Consistent with such a role, we found that SC differentiation was delayed in the absence of ZBTB20. In contrast, HC differentiation, as assessed by stereocilia morphology, remained unaffected in the absence of ZBTB20.

Likely the consequence of the ectopic proliferation of SC progenitors, we detected a mild but significant increase in the number of cochlear SCs in ZBTB20-deficient cochlear sensory epithelia at birth. In addition, analysis of ZBTB20-deficient cochlear sensory epithelia five days later revealed ectopic OHCs, a phenotype most pronounced at the apical tip. EdU pulses from P0-P5 failed to label cochlear HCs or SCs in *Zbtb20* KO mice, indicating that these extra SCs and HCs were not due to ectopic postnatal proliferation. Furthermore, ectopic OHCs in P5 *Zbtb20* KO mice had well-formed stereocilia and lacked SOX2 expression, suggesting they were formed at late embryonic stages. Our study and Xie et al. found that ZBTB20 loss arrests the development of Hensen cells (Xie et al., 2023), which may have led to their conversion into OHCs. Alternatively, it is possible that changes in planar cell polarity or mechanical forces due to defects in SC differentiation cause “bulging” of the epithelium and sporadic occurrences of four rows of HCs.

How does ZBTB20 regulate SC differentiation/maturation? Our transcriptomic data show that ZBTB20 loss delays the induction of the maturation-specific gene program. Importantly, it also revealed that the key stemness-associated genes *Sox11* and *Hmga2* were upregulated, suggesting that cochlear SCs were not only delayed in acquiring cell type-specific genes but also harbored an immature progenitor-like regulatory program that is largely decommissioned by this stage (P5) in wild-type tissue. This key finding provides explanatory insight into the response observed in regenerative assays with cochlear SCs from Emx2^Cre^ *Zbtb20* KO mice. Previous studies in our lab have shown that *Hmga2* is required for SC cell-cycle reentry and HC formation in organoid culture (Li et al., 2023), drawing a clear connection between the observed maturation defects of cochlear SCs in Emx2^Cre^ *Zbtb20* KO mice and their increased capacity to form organoids, proliferate, and produce HCs *in vitro*.

However, this immature cell state must not be the sole source of the increased SC regenerative capacity, as we also observed increased cell-cycle reentry and organoid formation in response to acute (TetO-Cre)-mediated deletion of *Zbtb20*. In this paradigm, we employed transcriptomic profiling of the expansion-phase organoids and identified the MDK-PTPRZ1 signaling axis as a potential target through which ZBTB20 may limit cochlear SC plasticity. We confirmed the efficacy of modulating this pathway by externally adding MDK, which also increased the proliferative and organoid-forming capacity of P5 cochlear SCs (and KCs). These results indicate that ZBTB20 in postnatal cochlear SCs continues to be required for reinforcing a terminal differentiated state.

A key unresolved issue is the differential impact of chronic versus acute ZBTB20 loss in cochlear tissue explants. We find that chronic ZBTB20 loss enables HC regeneration in both undamaged and damaged tissues, whereas acute deletion supports HC regeneration only in damaged tissues. The effectiveness of chronic ZBTB20 loss is likely attributable to delayed SC maturation and sustained expression of stemness-associated genes. In contrast, transcriptomic data suggest that acute ZBTB20 loss does not trigger de-differentiation or the acquisition of progenitor gene expression. HC loss at perinatal stages has been shown to trigger spontaneous HC regeneration (Bramhall et al., 2014; Cox et al., 2014). Recent findings indicate that HC damage during the perinatal period reactivates progenitor genes such as *Lin28b* and *Hmga2*, thereby enhancing cochlear SC plasticity (Li et al., 2022). Thus, pathways derepressed/activated by the acute loss of ZBTB20 may act synergistically with damage-induced pro-regenerative pathways to promote SC-to-HC transdifferentiation.

## METHODS

### Mouse Breeding and Genotyping

All procedures and experiments adhered to NIH-approved standards and were approved by the Johns Hopkins University Institutional Animal Care and Use Committee protocol. *Zbtb20* floxed mutant mice (RRID: MGI:5428606) were provided by Ulrich Mueller, Johns Hopkins University School of Medicine. *Emx2Cre/+* mice were obtained from Shinichi Aizawa, RIKEN (RRID: MGI: 3579416). *TetO-Cre* (RRID: IMSR_JAX:006234) and *R26rtTA*M2* (RRID: IMSR_JAX:006965) mice were obtained from Jackson Laboratories (Bar Harbor, ME). Both male and female mice were used in this study.

### RNA Extraction and RT-qPCR

Total RNA from organoids and tissue was extracted using the RNeasy Micro Kit (QIAGEN, no. 217084). Organoids were treated with Cell Recovery Solution (Corning, no.354253) prior to cell lysis to dissolve the Matrigel matrix. The iScript cDNA synthesis kit (Bio-Rad, no. 1708890) was used to reverse transcribe mRNA into complementary DNA (cDNA). QPCR was done on a CFX-Connect Real-Time PCR Detection System with SYBR Green Master Mix (Thermo Fisher Scientific, no. 4385612). *Rpl19* was used as an endogenous reference transcript. The ΔΔCT method was used to calculate relative gene expression.

### Cell Cycle Assays

For timed mating experiments, embryonic day E0.5 was defined as noon on the day a mating plug was observed. EdU (25 mg/kg, Thermo Fisher Scientific E10187) was administered via intraperitoneal injection to the pregnant dam on embryonic day E13.5, and pups were harvested just after birth at postnatal day P0. EdU incorporation was detected using Click-iT Edu Alexa Fluor 647 imaging Kit (Thermo Fisher Scientific, no. C10338).

### RNA Sequencing and Data Analysis

RNA was isolated from the isolated sensory epithelia of postnatal day 5 mouse cochleae of *Zbtb20^f/f^; Emx2^Cre/+^* (cKO), *Zbtb20^f/+^; Emx2^Cre/+^* (Het), and *Zbtb20^f/f^; Emx2^Cre/+^* (Ctrl) mice. RNA was extracted using the RNeasy Micro Kit, and samples were subsequently processed using Illumina’s TruSeq stranded Total RNA kit according to manufacturer recommendations and sequenced using the Illumina NovaSeq 6000. For RNA-seq experiments in organoid culture, Ctrl (T*etO^+/+^; R26^M2rtTA^; Zbtb20^f/f^*) and KO (*TetO^Cre/+^; R26^M2rtTA^; Zbtb20f/f) cultures were treated with Cell Recovery solution,* followed by RNA extraction. These samples were shipped on dry ice and sequenced by Plasmidsaurus using the Illumina NovaSeq X Plus. All reads were pseudo-aligned to a reference transcriptome (Ensembl v96) using kallisto (v0.46.1), and differential testing was performed using sleuth (v0.30.2). Gene Ontology was performed using Metascape (v3.5) and clusterProfiler (v4.18.4), and graphics were constructed using ggplot2 (v4.0.2).

### Immunohistochemistry

Dissected cochleae were fixed using 4% (vol/vol) paraformaldehyde in Phosphate Buffered Saline (PBS) (Electron Microscopy Sciences, no. 15710) for at least 12 hours at 4°C. Cell membrane permeabilization was achieved by incubation in 0.1% (vol/vol) TritonX-100 in PBS for 3 times, 10 minutes each, at room temperature. Then, samples were incubated in a blocking buffer of 0.5% (vol/vol) TritonX-100, 5% (vol/vol) Donkey Serum (Millipore Sigma, D9663), and 10% Bovine Serum Albumin (Millipore Sigma, A1595) in PBS for 1 hour at room temperature. Primary antibodies were added to the blocking solution, and samples were incubated for at least 12 hours at 4°C. After washing with 0.1% (vol/vol) TritonX-100 in PBS 3 times for 10 minutes each at room temperature, samples were incubated with secondary antibodies and, for some experiments, phalloidin 488 (Thermo Fisher Scientific, A12379) and DAPI (Biolegend, 422801) stains for at least 2 hours at room temperature. Finally, samples were mounted onto microscope slides (Fisher, 12-550-15 using Fluoromount G (Invitrogen, 00-4958-02. For samples requiring EdU labeling, labeling was performed before adding secondary antibodies.

### Statistical Analysis

Animals (biological replicates) were assigned to experimental or control groups based on genotype and/or treatment. Sample size (n) represents the number of animals that were analyzed per genotype/treatment. Data were analyzed using R (4.5.3) and GraphPad Prism (11.0.2), and masking was employed to avoid bias. Relevant information for each experiment, including the sample size analyzed, the statistical tests applied, and the p-values reported, is provided in the corresponding figure legend. In most cases, p-values ≤ 0.05 were considered significant, with the exception of RNAseq analysis, for which a q-value ≤ 0.05 was used to correct for multiple comparisons. On all graphs, error bars represent standard deviations.

### Organoid culture

Cochlear sensory epithelia were enzymatically separated from surrounding tissue, incubated in TrypLE (Thermo Fisher Scientific, no. 2604013) and triturated to dissociate to a single-cell level, and plated in a drop of Matrigel matrix at high density (3000 cells, stage P5; 4000 cells, stage P7). The droplets were cultured in a pro-growth medium [DMEM/F12 (Corning, no. 10–092-CV), 1X N-2 (Thermo Fisher Scientific, no.17502048), 1X B27 (Thermo Fisher Scientific, no.12587010) and penicillin (100 U/ml; Sigma-Aldrich, no. P3032)] supplemented with EGF (5 ng/ml; Sigma-Aldrich, no. SRP3196), FGF2 (50 ng/ml; Thermo Fisher Scientific, no. PHG0264), CHIR99021 (3 μM; Sigma-Aldrich, no. SML1046), 616 (Sigma-Aldrich, no. 61654), and VPA (1 mM; Sigma-Aldrich, no. P4543). To induce differentiation, the culture medium was supplemented with CHIR99021 (3 µM) and LY411575 (5 µM; Sigma-Aldrich, no. SML0506). For genetic models employing an inducible Cre recombination mechanism, doxycycline hyclate (0.5 µg/ml) was added to the expansion medium.

### Fluorescence-activated cell sorting

For postnatal day 7 mice, cells from cochlear epithelia were collected and dissociated as previously described (*citation*). For postnatal day 13 mice, whole cochleae for each animal were collected and dissociated. Cells were then suspended in Hanks Balanced Salt Solution (minus CaCl_2_ and MgCl_2_) (Thermo Fisher Scientific 14175095) containing 2% (vol/vol) Fetal Bovine Serum (Gibco, 26-140-079), incubated with propidium iodide (PI), and sorted on a MoFlo Legacy sorter with a 100-μm nozzle tip. Lfng-GFP-positive, PI-negative cells were collected in organoid expansion medium and cultured as described above.

### Quantification of organoid formation efficiency and organoid diameter

Low-power bright-field images of organoids were captured with an Axiovert200 microscope using 2.5X, 5X, and 10X objectives (Carl Zeiss Microscopy). Organoid formation efficiency was measured by counting organoids at 10 days in vitro and dividing by the total number of cells plated. The total number of plated cells was established at the time of culture generation using a cell counter (BioRad TC 20 Automated Cell Counter). Diameters were measured in FIJI (2.14.0). For each genotype and/or condition, at least three different animals per independent experiment were included in the analysis.

### Explant culture

Organotypic cochlear explants, which are comprised of the cochlear sensory epithelia and innervating neurons, were microdissected from stage P2 mice and cultured on SPI-Pore membrane filters (Structure Probe, no. E1013-MB) in DMEM/F12 containing 1X N-2, 1X B27, EGF (5 ng/ml), CHIR99021 (3 μM, and penicillin (100 U/ml). To ablate HCs, diphtheria toxin (25ng/mL, Millipore Sigma, D0564-1MG) was added to cultures for 24 h, followed by a washout. Cultures were supplemented with LY411575 (5 μM) to induce HC formation. For genetic models employing an inducible Cre recombination mechanism, doxycycline hyclate (0.5 µg/ml) was added to the explant culture medium.

## ACKNOWLEDGEMENTS

We thank the members of the Doetzlhofer Laboratory for their help and advice throughout this study.

## AUTHOR CONTRIBUTIONS

Conceptualization: A.D., C.M. Methodology: A.D., C.M., Z.U.R. Investigation: C.M., Z.U.R. Supervision: A.D. Writing: A.D., C.M.

## DATA AVAILABILITY

RNA sequencing data have been deposited in the Gene Expression Omnibus data repository under accession GSE342694.

