## Supplemental Figures for "Loss of Zbtb20 disrupts cochlear supporting cell differentiation and maturation and extends the postnatal hair cell regenerative window in mice"

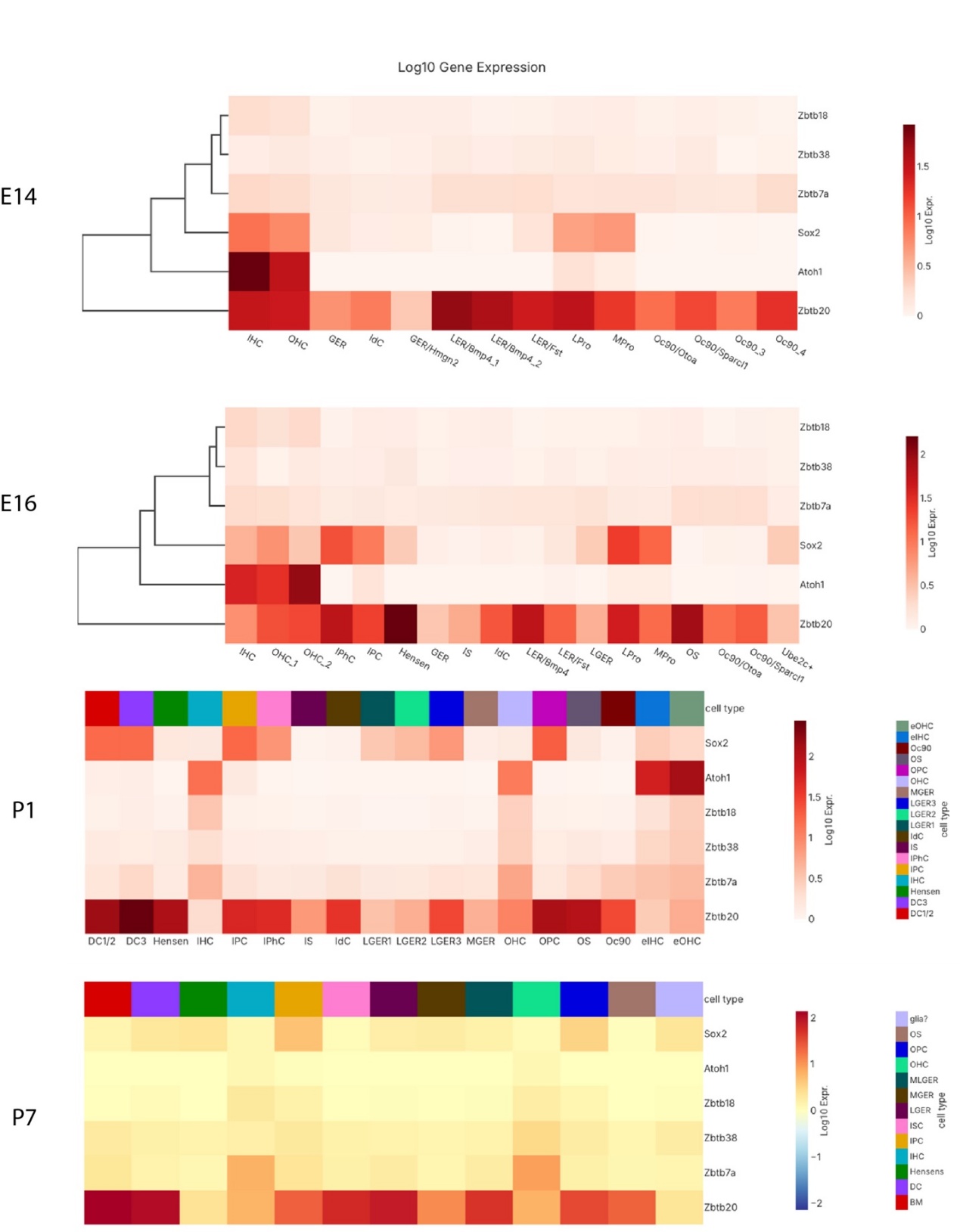

**Fig. S1.** *Zbtb20* expression in the embryonic and early postnatal cochlea.

(A) Heatmaps showing the expression patterns of *Zbtb20* and other genes in the ZBTB family of transcription factors in the mouse cochlea at E14, E16, P1, and P7. *Atoh1* and *Sox2* are included to denote SC and HC subtypes

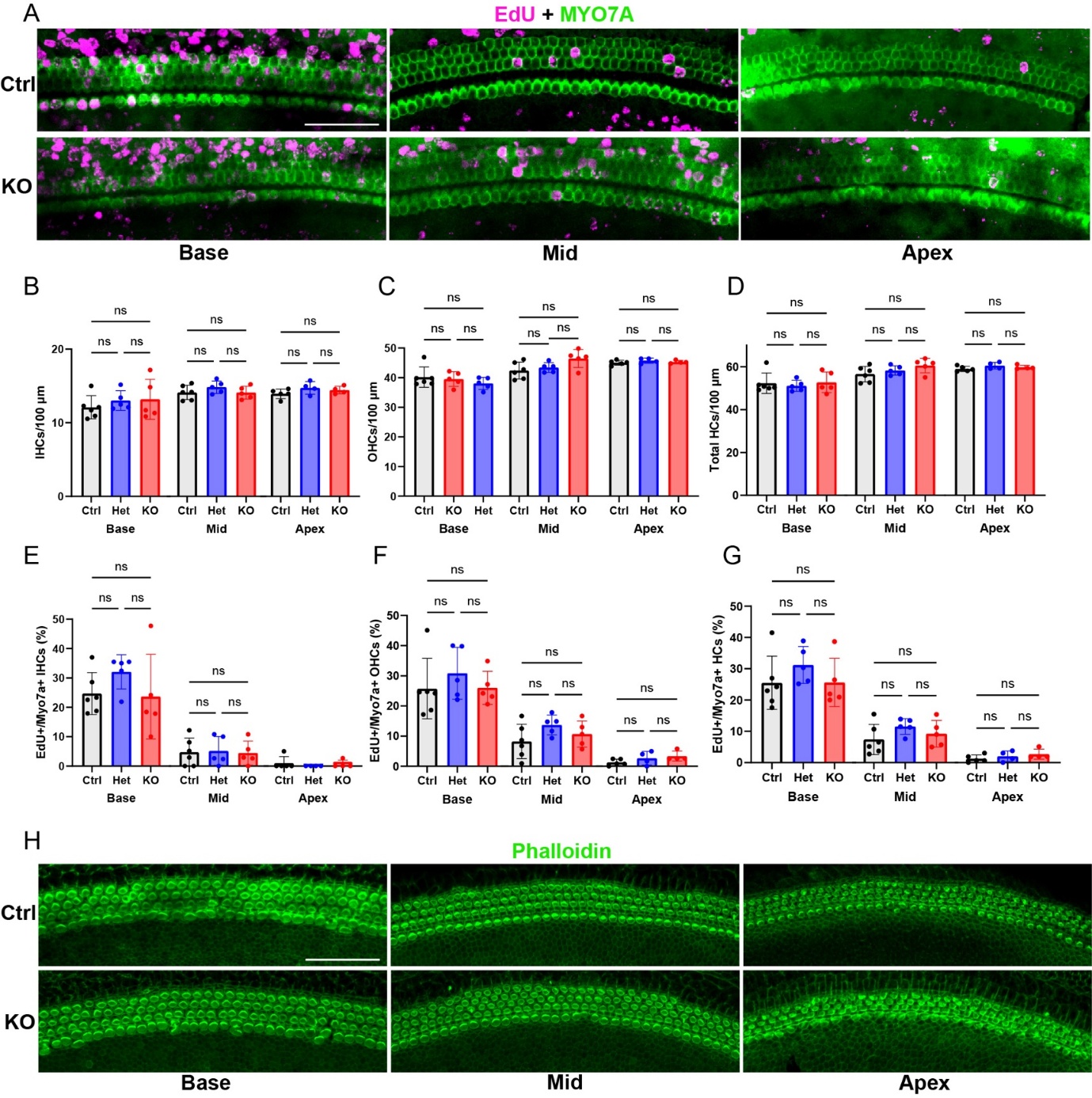

**Fig. S2.** ZBTB20 loss does not delay cell cycle exit or differentiation of cochlear HC progenitors.

(A) Representative confocal images of EdU incorporation (magenta) in HCs (MYO7A, green) at the base, mid, and apex of cochlear sensory epithelia from P0 *Emx2^Cre^ Zbtb20* KO mice and control littermates. Timed-pregnant female mice received a single EdU pulse at E13.5 as in Fig. 1D. Control, het, and KO mice were collected. Scale bar = 50μm.

(B) Quantification of inner hair cells in (A). *P* values were determined using a one-way ANOVA and post-hoc Tukey’s HSD test for multiple comparisons.

(C) Quantification of outer HCs in (A). *P* values were determined using a one-way ANOVA and post-hoc Tukey’s HSD test for multiple comparisons.

(D) Quantification of all HCs in (A). *P* values were determined using a one-way ANOVA and post-hoc Tukey’s HSD test for multiple comparisons.

(E) Quantification of EdU incorporation in IHCs as shown in (A). *P* values were determined using a one-way ANOVA and post-hoc Tukey’s HSD test for multiple comparisons.

(F) Quantification of EdU incorporation in OHCs as shown in (A). *P* values were determined using a one-way ANOVA and post-hoc Tukey’s HSD test for multiple comparisons.

(G) Quantification of EdU incorporation in all HCs as shown in (A). *P* values were determined using a one-way ANOVA and post-hoc Tukey’s HSD test for multiple comparisons.

(H) Representative confocal images showing phalloidin labeling of F-actin (green) to show HC stereocilia at the base, mid, and apex of P0 control and KO cochleae. Scale bar = 50μm.

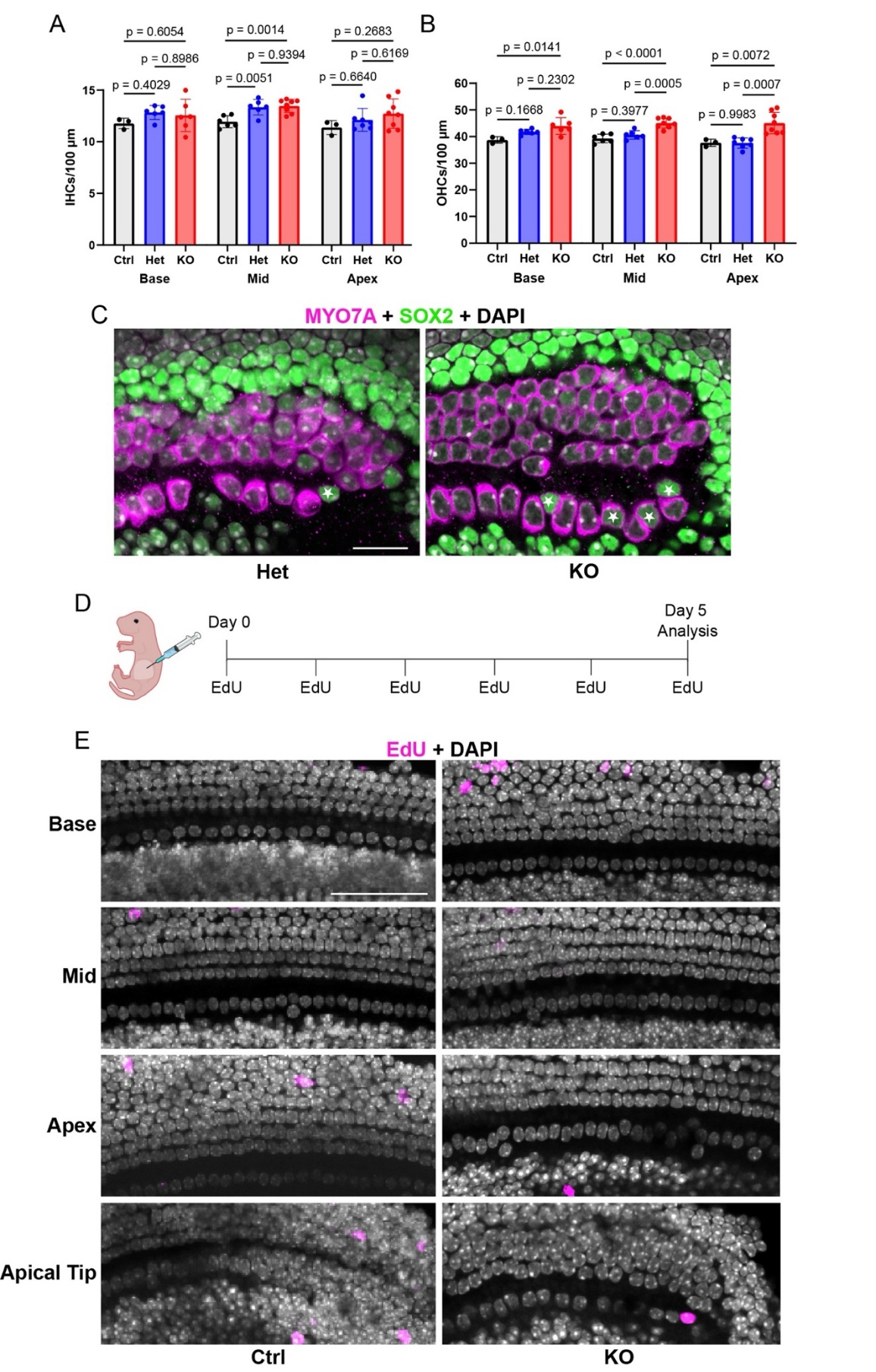

**Fig. S3.** Loss of *Zbtb20* does not induce ectopic proliferation in the cochlear sensory epithelium at early postnatal stages.

(A) Quantification of IHCs in cochlear sensory epithelia of P0 *Emx2^Cre^ Zbtb20* KO mice, het and control littermates as shown in Fig. 2A. *P* values were determined using a one-way ANOVA and post-hoc Tukey’s HSD test for multiple comparisons.

(B) Quantification of OHCs in Fig. 2A. *P* values were determined using a one-way ANOVA and post-hoc Tukey’s HSD test for multiple comparisons.

(C) Representative confocal images of the cochlear apical tip in het and KO mice. Stars are placed to highlight SOX2 labeling present in IHCs. Scale bar = 20μm.

(D) Schematic of EdU injections and tissue harvesting.

(E) Representative confocal images of EdU labeling at the base, mid, and apex of P5 control and KO cochleae. Scale bar = 50μm.

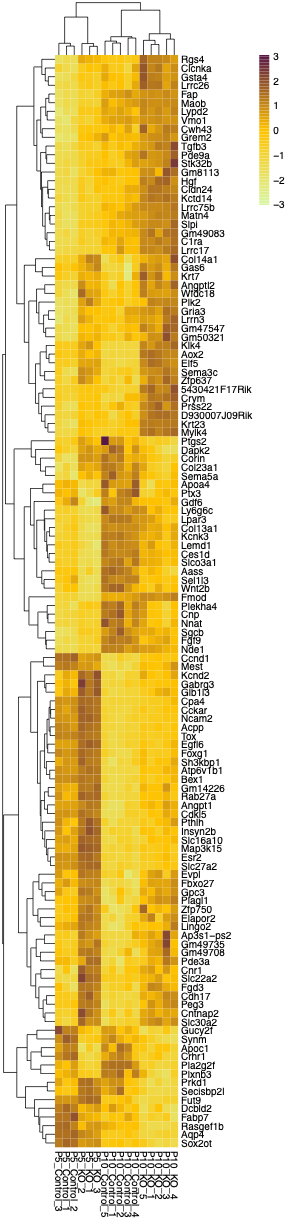

**Fig. S4.** Full annotated heatmap of genes that were differentially expressed in both P5 sensory epithelia (control vs KO) and P10 whole cochlea (control vs KO) as shown in Fig. 3F.

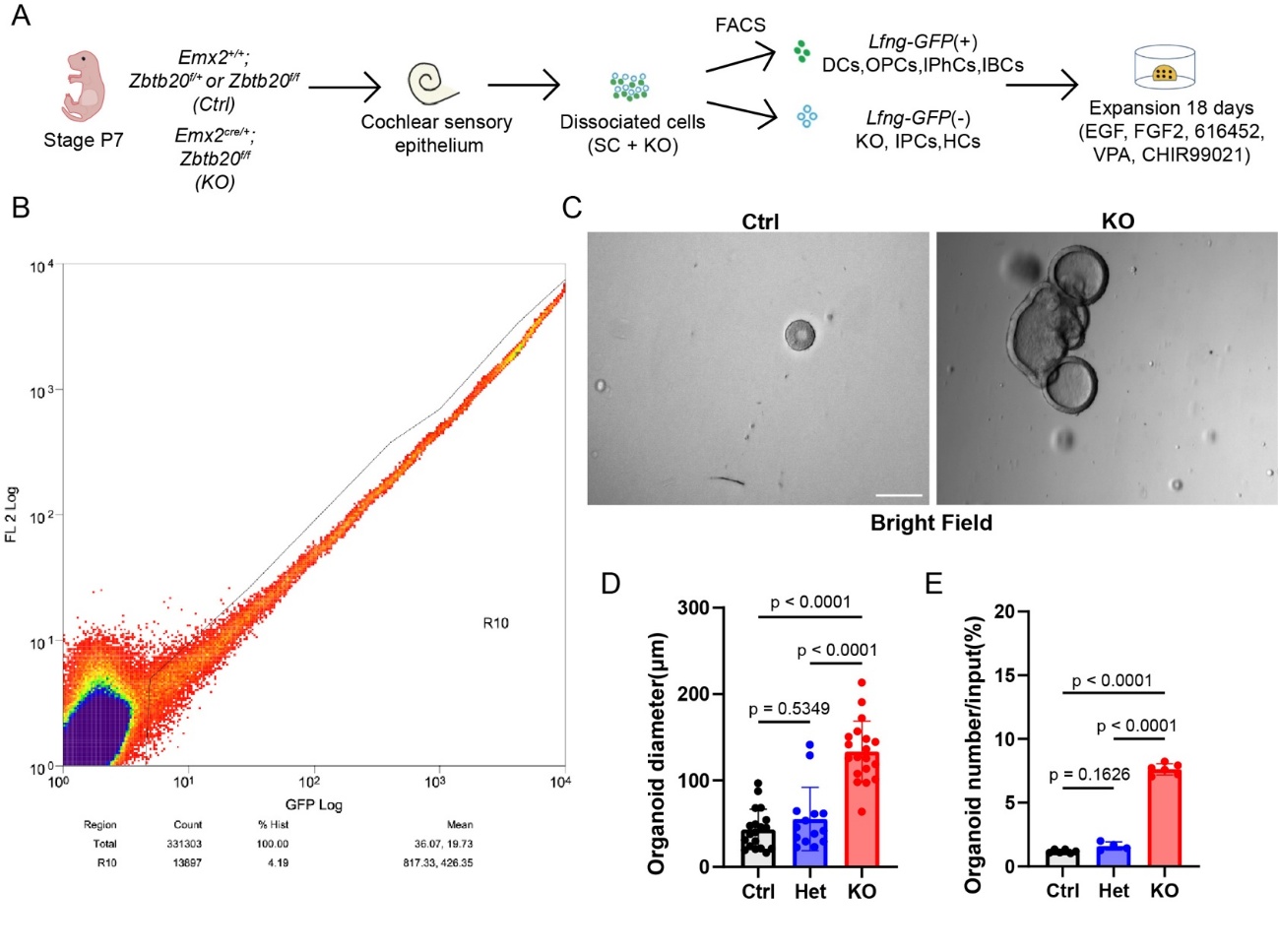

**Fig. S5.** Chronic deletion of *Zbtb20* enhances the proliferative and organoid-forming capacity of P7 cochlear *Lfng-GFP*(+) supporting cells.

(A) Experimental schematic of organoid generation and HC differentiation. Fluorescence-activated cell sorting (FACS) was used to enrich for *Lfng-GFP*(+) Deiter cells (DCs), outer pillar cells (OPCs), inner phalangeal cells (IPhCs), and inner border cells (IBCs) as well as to remove dead cells. Made with graphics from scidraw.io.

(B) Representative graph showing gating parameters used to enrich for *Lfng-GFP*(+) cells during FACS.

(C) Representative bright-field images of control and KO organoid cultures after 18DIV. Scale bar = 100μm.

(D) Quantification of organoid diameter as shown in (C). *P* values were determined using a one-way ANOVA and post-hoc Tukey’s HSD test for multiple comparisons.

(E) Quantification of organoid formation efficiency as shown in (C). *P* values were determined using a one-way ANOVA and post-hoc Tukey’s HSD test for multiple comparisons.
